# How copper based grape pest management has impacted wine aroma

**DOI:** 10.1101/2023.11.12.566766

**Authors:** Irene De Guidi, Virginie Galeote, Bruno Blondin, Jean-Luc Legras

## Abstract

Despite the high energetic cost of the reduction of sulfate to H_2_S, required for the synthesis of sulfur-containing amino acids, some wine *Saccharomyces cerevisiae* strains have been reported to produce excessive amounts of H_2_S during alcoholic fermentation, which is detrimental to wine quality. Surprisingly, in the presence of sulfite, used as a preservative, wine strains produce more H_2_S than wild (oak) or wine velum (*flor*) isolates during fermentation. Since copper resistance caused by the amplification of the sulfur rich protein Cup1p is a specific adaptation trait of wine strains, we analyzed the link between copper resistance mechanism and H_2_S production. We show that a higher content of copper in the must increases the production of H_2_S. Using a set of 51 strains we observed a positive and then negative relation between the number of copies of *CUP1* and H_2_S production during fermentation. This complex pattern could be mimicked using a multicopy plasmid carrying *CUP1*, confirming the relation between copper resistance and H_2_S production. The massive use of copper for vine sanitary management has led to the selection of resistant strains at the cost of a metabolic tradeoff: the overproduction of H_2_S, resulting in a decrease in wine quality.

## Introduction

The most ancient traces of wine making have been discovered in Georgia ^1^ and have been dated as 6000 BC. Since that ancient time, cultivation of grapevine and winemaking knowledge spread progressively all over the world ^2^. All along this period, winemaking practices have evolved, especially with the discovery of the use of sulfite to limit the growth of undesired microorganisms to protect wine from oxygen and to preserve aroma profile. Similarly, the cultivation of *Vitis vinifera* has faced changes, especially with the development of grafting and the spray of chemical compounds required to face the import in Europe of three major pests for vine: phylloxera, powdery mildew and downy mildew. Among chemicals sprayed on vines, copper has been intensively used in vineyards to control the development of *Plasmopara viticola*. This intensive use of copper in vineyards has translated into high copper in grape musts ^3^.

Wine fermentation is mainly achieved by the yeast species *Saccharomyces cerevisiae* which is found also in many fermented products: sake, bread, cheese and more ^4–7^. This yeast species can also be encountered in natural biotopes such as forests ^8,9^. *S. cerevisiae* strains display specific physiological properties associated to the different ecological niches they live in, as result of several domestication events^4,10–12^.

One of the most remarkable and contrasting adaptation events can be seen in fermenting wine strains and in wine velum isolates (*flor* yeasts). *S. cerevisiae* velum strains have developed a specialized aerobic lifestyle, highly different from the one of fermenting wine strains ^13^. Since they colonize the wine when fermentation is concluded, velum strains develop the ability to grow in media depleted for nitrogen, vitamins, glucose and fructose. The entire wine fermentation process constitutes a difficult environment *S. cerevisiae* has to cope with, and different features of the genome of wine strains have been recognized as traces of adaptation associated with their domestication ^14^. The first and best described adaptation of *S. cerevisiae* to the wine environment is the resistance to sulfite, obtained from several translocation events resulting in a high expression of the sulfite export pump Ssu1 ^15–19^. Another example of adaptation to the grape and must environments can be seen in the selection of strains carrying multiple copies of the *CUP1* gene. This gene amplification leads to an enhanced protein abundance/synthesis, providing resistance to high concentrations of copper in the grape must, resulting from the massive use of copper as fungicide ^20^. Cup1p is among the ten sulfur richest yeast proteins ^21^ and some *S. cerevisiae* strains can harbor up to 79 copies ^18,20,22^. Therefore, the high synthesis of Cup1p caused by its amplification requires a high availability of sulfur containing amino acids methionine and cysteine that are scarce in grape musts. These amino acids can be synthesized by yeast through the sulfur assimilation pathway (SAP), which reduces inorganic sulfate into hydrogen sulfide (H_2_S) with the consumption of 7 moles of NADPH and 4 of ATP per mole of S-amino acid ^23^. Consequently, the biosynthesis of the sulfur amino acids has a significant impact on the yeast redox and energy balances. A high diversity in the production of H_2_S during alcoholic fermentation has been described for wine strains ^24^, and because its content is detrimental to wine aroma, different studies have deciphered its genetic bases and found allelic variations in *MET10, SKP2, MET2, TUM1* ^25–28^, genes involved in the sulfur assimilation pathway or its regulation. Some of these findings have been patented and have led to the improvement of industrial winemaking starters. Surprisingly, no investigation has been carried out to understand the biological meaning of such overproduction, nor to evaluate a potential relation with different ecological niches. Interestingly, for wine *S. cerevisiae*, SO_2_ and copper tolerance have been found negatively associated ^29^. Transcriptional and proteomic analysis in sulfur-limited medium, demonstrated that *SSU1* over-expression induced sulfur limitation during exposure to copper and provoked an increased sensitivity to copper ^30^.

Because the production of H_2_S is so costly to the cell ^23^, we wondered why some wine strains were overproducing it. Comparing three groups of strains: isolated from velum and wine, two contrasted anthropogenic environments, and oak, as a natural environment, we show here that the total content of H_2_S produced during alcoholic fermentation depends on the ecological niche, and that exposure of yeast cells to copper enhances H_2_S production. Last, evaluate how the amplification of *CUP1* may explain such variation, using a set of strains with variable number of copies of *CUP1* or strains carrying a plasmid overproducing *CUP1*.

## Results

### Strain variability in H_2_S production during alcoholic fermentation

To assess the variability of the production of H_2_S during alcoholic fermentation of *Saccharomyces cerevisiae*, we evaluated 33 strains isolated from three ecological niches: wine (n=10), wine velum (n=14), and from oak trees (n=9), as a wild reference. Because SO_2_ is an intermediate of the sulfur assimilation pathway, and used in most wine fermentations as an additive for its antimicrobial, antioxidant and anti-oxidizing activities, we compared the H_2_S production of the mentioned ecological groups in a synthetic grape must in absence or presence of sulfites. The variability in H_2_S production among strains of these three groups is presented in Figure 1 and Supplementary Figure 1. A two-way ANOVA revealed that the origin of the strain has a significant effect on H_2_S produced (F_2,128_=36.31, p_value=3.24x10^-13^) as well as the addition of SO_2_ to the must (F_1,128_=59.19, p_value=3.36x10^-12^). A significant interaction between the effects of the two factors (i.e. SO_2_ and origin) on H_2_S produced during alcoholic fermentation (F_2,128_=14.5, p_value=2.20x10^-6^) was detected.

**Figure 1:**
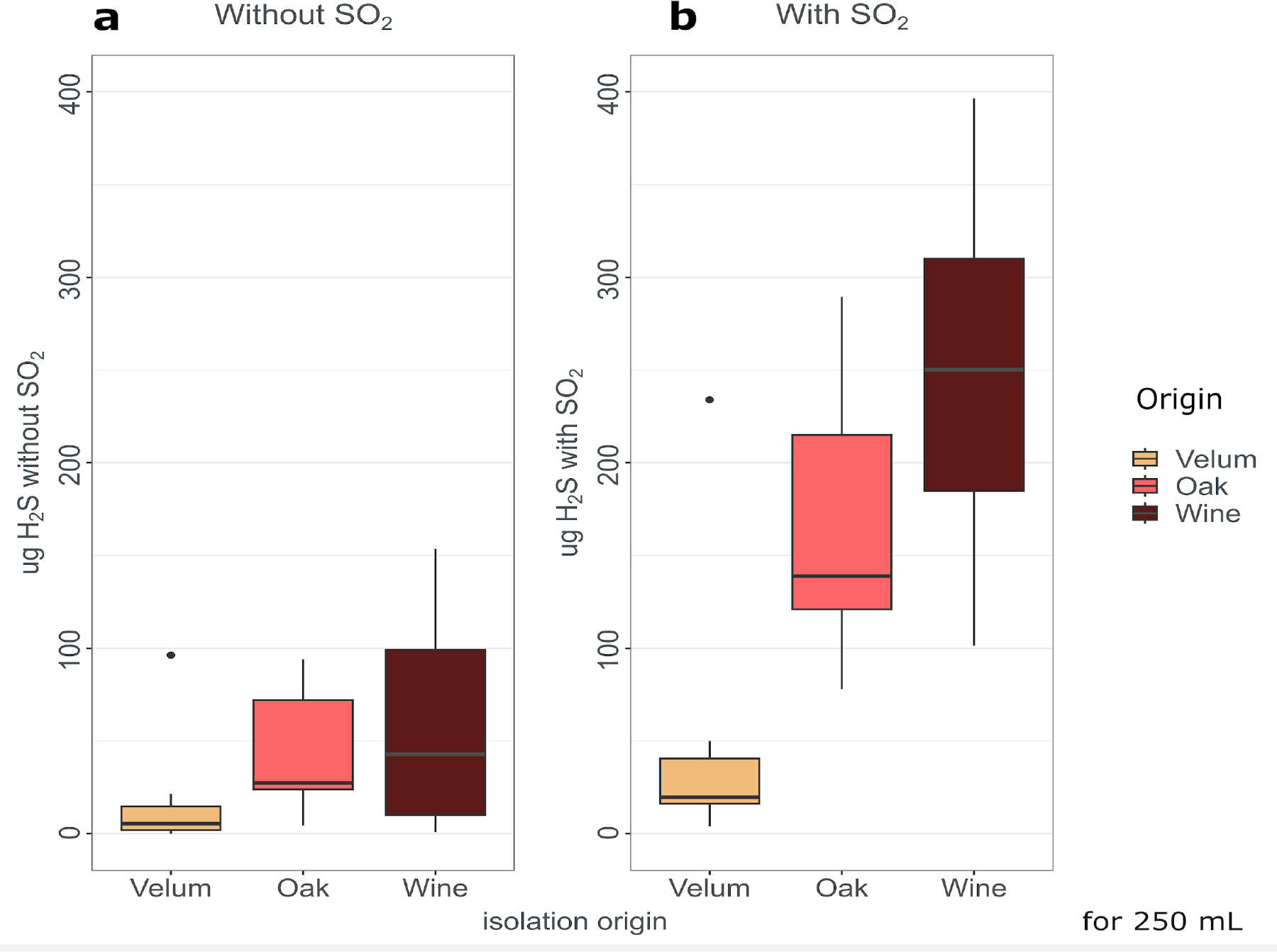
Difference in the average of cumulate H_2_S production during alcoholic fermentation of velum, oak and wine strains in absence (a) or presence (b) of SO_2_.

Tukey multiple comparisons of means at 95% family-wise confidence level showed that H_2_S production between the independent origins was significant when the must contained sulfite (Figure 1b). Wine strains produced more H_2_S compared to velum and oak (p_value=1.50x10^-6^ and 0.014 respectively). Oak isolates also produced more H_2_S than velum strains in the presence of sulfite (p_value=1.50x10^-6^). This difference was not noticeable when the must did not contain sulfites (Figure 1a).

Comparing figure 1a and 1b, it was clear that strains isolated from different origins did not respond with the same amplitude to the sulfite treatment. Velum strains displayed a remarkable low H_2_S production even in the presence of sulfite, in comparison to wine and oak strains (p_value=1.50x10^-6^ and 3.34x10^-5^ respectively). This explains the significant interaction detected by the model.

### Influence of copper content on H_2_S production

The copper content of the grape must or wine can result from the traces left with sanitary treatment performed on vine, and also from treatments aimed at reducing H_2_S production. Indeed, thiols functions reduce Cu^2+^ and produce a Cu^+^ that binds to -SH functions ^31^. We evaluated the effect of copper concentrations of the must on H_2_S production of two industrial winemaking starters: VL1, a wine strain, identified as low H_2_S producer in the first experiment, and LMD17, a high H_2_S producer wine strain (De Guidi et al. 2021)”. The analysis of variance revealed a significant effect of both factors: strain (F_1,14_= 57, p_value=2.7x10^-6^), and copper (F_2,14_=10.6, p_value=0.002). The results presented in Figure 2 show a clear increase of H_2_S production with an increase in copper content in the synthetic grape must at concentrations compatible with those encountered in winemaking ^3^. For both strains, higher concentrations provoke the formation of a black precipitate that hampers H_2_S measurement with our method, and suggest higher production (Supplementary Figure 2).

**Figure 2.**
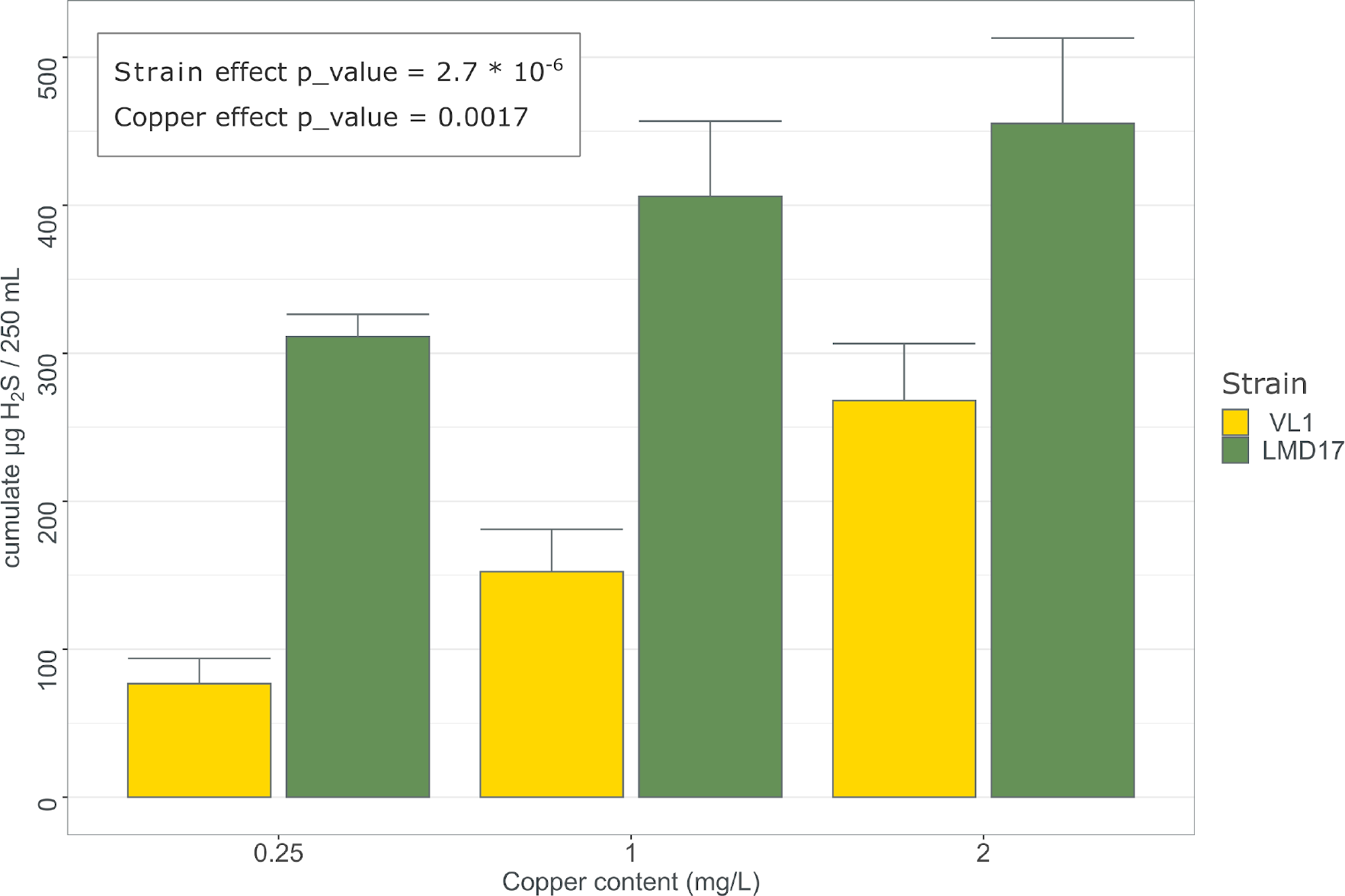
Effect of copper content in synthetic must without SO_2_ on total H_2_S production during alcoholic fermentation by wine strains VL1 and LMD17. p values refer to two-way ANOVA.

### Evaluating the relation between H_2_S production and *CUP1* copy number

Besides inducing H_2_S production, the copper content in the growth medium controls the expression of *CUP1* ^32^ that is involved in its detoxification. In addition, we observed that *CUP1* is one of the proteins with the highest sulfur containing aminoacid content (21.31%), just after *MNC1* (25.76%), another membrane protein that is upregulated by toxic concentrations of heavy metal ions ^33^.

In a first approach aimed at exploring the effect of *CUP1* copy number on H_2_S production in strains of the same three niches analysed above, we increased the number of strains to test (+18 wine isolates, total n = 51), in order to include strains with 2 to 71 *CUP1* copy number. Surprisingly, we observed a non-linear relation between t*CUP1* copy number and total H_2_S production.

As shown in Figure 3, strains with 1 to 10 *CUP1* copies exhibit an increasing total H_2_S production, whereas for more copies, it progressively decreases until it reaches almost null values. A 3^rd^ degree polynomial model described well the H_2_S production in relation to the *CUP1* copy number of the strains (black line in Figure 3), displaying a bell shape, that remained even after the removal of the two highest values for H_2_S production (red line in Figure 3). One possibility of this shape could result from an increasing activation of SAP to support *CUP1* production when up to at least 10 gene copies are present, whereas for higher number of copies, the higher requirement of sulfur amino acids might overcome the SAP maximum activity, leading to an increased utilization of H_2_S for the synthesis and a lower release. This increase in H_2_S concentration provoked by the increase number of copies of *CUP1* within the range 1-10 copies is similar to the effect caused by the copper content of the grape must observed for VL1 and LMD17.

**Figure 3.**
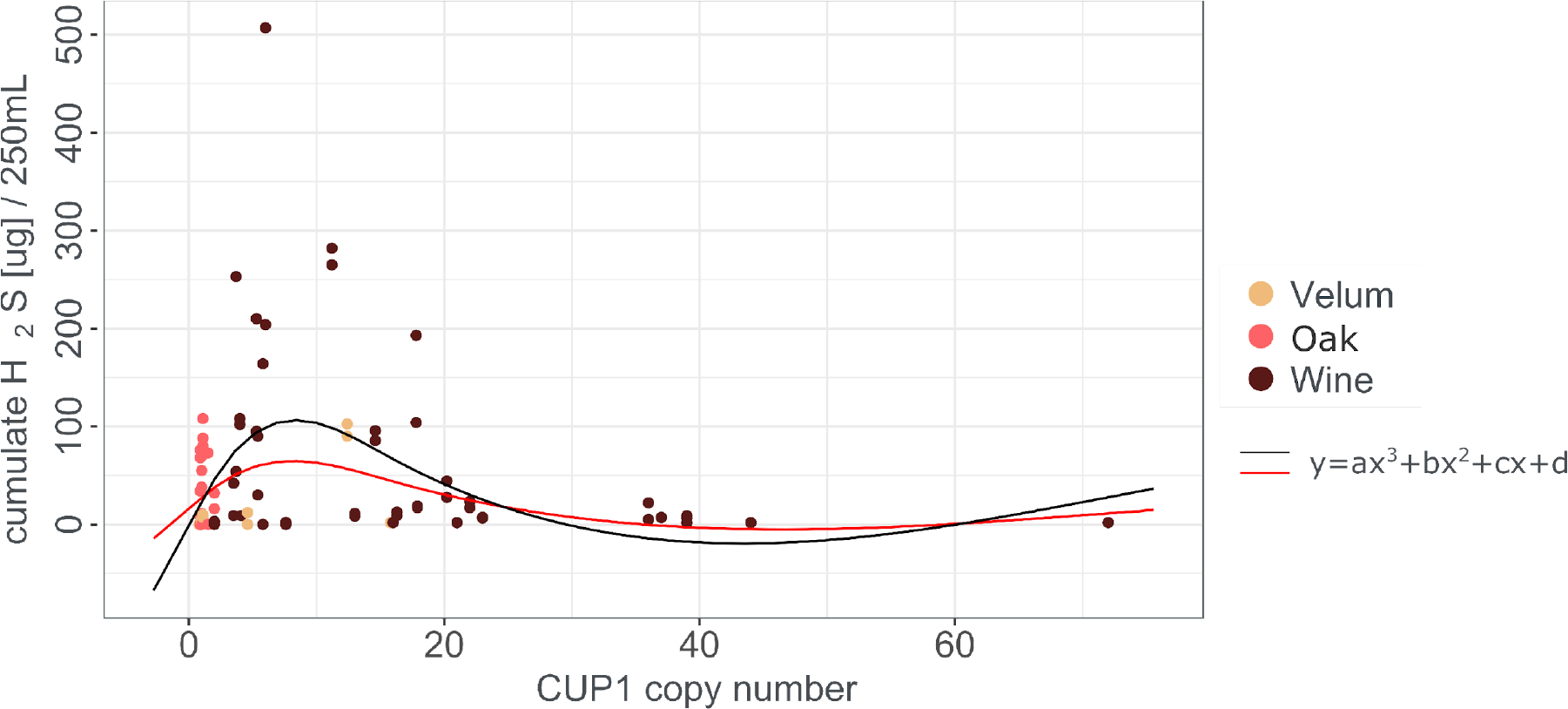
H_2_S production (in synthetic must without SO_2_) distribution as function of *CUP1* copy number of each strain. Different isolation origins (the same of Figure 1) are described by colors. Black solid line: 3^rd^ degree polynomial model describing the relation between H_2_S production and *CUP1* copy number; red line: same model excluding the two highest H_2_S producers.

### Impact of the modulation of *CUP1* copy number on H_2_S production

In order to validate the effect of *CUP1* copy number on H_2_S production, we tried to manipulate the number of *CUP1* copies per cell. With this aim, we built a multicopy yeast episomal plasmid (YEp) expressing *CUP1* under the control of the strong promoter from the translational elongation factor EF-1 alpha (*TEF1*). Three strains with different number of genomic copies of *CUP1* were transformed with this plasmid and tested in a media containing a low copper concentration (0.25 mg/L).

First, the overexpression of *CUP1* in the oak strain OAK-Rom 3_2, a low H_2_S producer with one copy of *CUP1*, led to a significant increase of H_2_S production (F_2,6_=9.61 p_value=0.013, Figure 4). In contrast to the oak strain, the overexpression of *CUP1* in the wine strain LMD17, ahigh H_2_S producer, with 11 copies of *CUP1*, decreased H_2_S production by half (F_2,5_=17.49, p_value=0.006, Figure 4). Last, the overexpression of *CUP1* in L1374, which carries 36 copies of *CUP1* and was ranked among the lowest H_2_S producers, did not change its production (F_2,6_=0.2, p_value=0.824, Figure 4). The responses displayed by these three constructions are in agreement to the experimental data presented in Figure 3, reproducing the “bell-shape” trend of H_2_S production.

**Figure 4.**
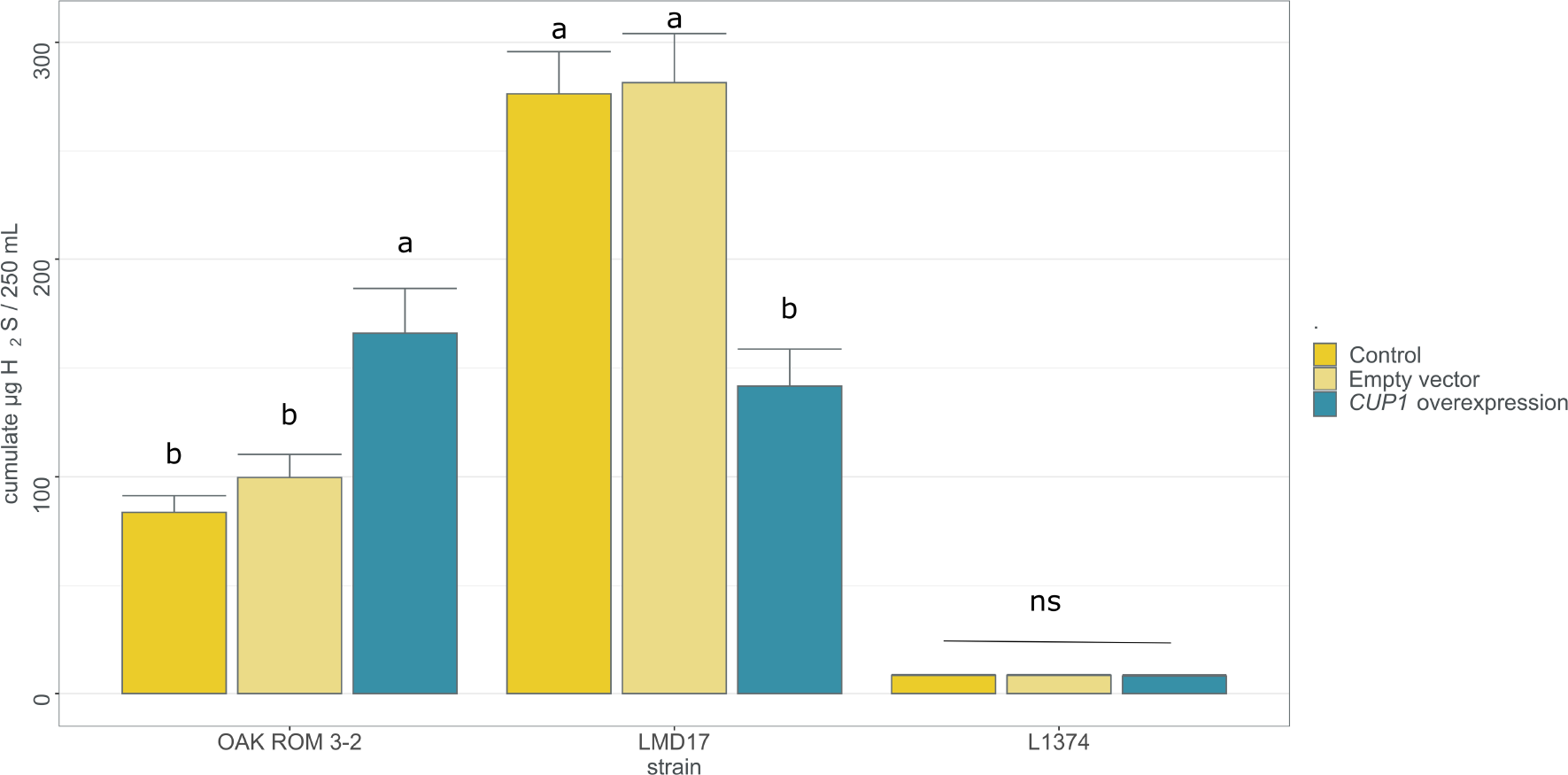
Effect of *CUP1* overexpression on total H_2_S released during alcoholic fermentation (in synthetic must without SO_2_) in three *S. cerevisiae* strains. Different lowercase letters indicate statistically significant differences between the molecular modifications (control wild-type strain, empty vector or *CUP1* overexpressing vector) for each strain separately, after Tukey multiple comparison of means at 95% family-wise confidence level.

## Discussion

Our results, obtained under conventional winemaking conditions (i.e., in the presence of sulfites), demonstrate for the first time that the production of H_2_S by *Saccharomyces cerevisiae* during alcoholic fermentation varies among natural and two distinct groups of domesticated strains. Surprisingly, we observed that wine populations exhibit the highest H_2_S production when sulfites are added to the grape must, which is unexpected considering that H_2_S is highly undesirable in winemaking. The increase in H_2_S production induced by sulfite addition is understandable as this compound is an intermediate metabolite in the sulfur assimilation pathway. However, it was surprising to find significant differences between wine and oak strains, whereas wine strains carry several types of translocations leading to a higher expression of the sulfite efflux pump *SSU1*. This contradicts the antagonistic role of *SSU1* in copper resistance ^30^, suggesting a limitation in the sulfur assimilation pathway. However, it should be noted that Onetto *et al*.’s study was conducted in the absence of added sulfites. We can hypothesize that the addition of sulfites to the grape must, as performed by winemakers, overflows the sulfite export transporter’s expulsion capacity, resulting in a higher production of H_2_S in wine strains.

We also show that the presence of copper in the grape must enhances the production of H_2_S. Our results align with the expression data obtained previously, which demonstrate the upregulation of genes encoding the two subunits of the sulfite reductase *MET5, MET10* (i.e., the main enzyme of the sulfur assimilation pathway), and the protein responsible for copper resistance/detoxification *CUP1* following copper exposure ^34,35^. Furthermore, the metallothionein protein Cup1p, which has one of the highest contents of sulfur-containing amino acids methionine and cysteine in the *S. cerevisiae* proteome, requires the availability of these amino acids for its synthesis. The differences in H_2_S production among oak, wine, and velum yeast strains may be attributed to variations in the number of *CUP1* copies in their genomes. However, we describe a complex relationship between the number of *CUP1* copies and H_2_S production. Moderate amplification of *CUP1* (up to approximately 10 copies) leads to an increase in H_2_S production, whereas higher copy numbers result in a lower fraction of H_2_S being stripped by the CO_2_ generated during fermentation. In this case, we propose that this reflects a higher utilization of H_2_S for amino acid synthesis.

Lastly, our results also highlight the specific behavior of flor strains. Unlike wine strains, velum strains exhibit very low H_2_S production, and have a lower number of *CUP1* copies. Velum strains grow at the surface of wine, after alcoholic fermentation which has significantly reduced the copper content of wine. It is likely that the selection pressure for copper-resistant strains has been less intense for flor strains compared to wine yeast. Another possible explanation for the lower H_2_S production in velum strains is the reduced activity of the pentose phosphate pathway, which provides NADPH, in comparison to oak breads and wine strains ^36^. This sheds light on the different domestication trajectories of wine and flor strains, reflecting their distinct lifestyles ^13^.

## Conclusion

The long-term exposure of yeast to copper, used for vine pest management over 150 years, has promoted their adaptation by selecting strains with multiple copies of *CUP1*. Our results indicate that this adaptation comes with a significant trade-off: increased resistance to copper used for pest management but also a higher production of H_2_S by the yeast, which has a detrimental impact on wine quality. This elevated H_2_S production is further amplified in the presence of sulfite, another additive commonly used in winemaking. Given the energetic cost of the H_2_S production, the impact on the global yeast metabolism should be evaluated. Numerous projects and techniques have been launched to mitigate or limit H_2_S production ^26,27,37^ and despite significant advances in the understanding of the production of this unpleasant gas, none of them investigated how the use of copper may have caused this phenotype. The yeast *CUP1* background should be considered when looking for the selection of wine yeast for low H_2_S production. Nevertheless, diversity data suggests that the amplification of *CUP1* is very likely not the sole mechanism explaining variations in H_2_S production, and more investigations have to be performed.

## Materials and Methods

### Strains

Fifty-one *Saccharomyces cerevisiae* from different geographical areas were characterized for their H_2_S production during alcoholic fermentation. The genetic group, identified in previous works indicated in the references of Supplementary table 1, reflected the colonized ecological niche: 28 belong to the “wine” clade, 14 to “velum” group and 9 to the “oak” one. Strains were selected from our laboratory collection and maintained on solid medium (agar YPD: 2% glucose, 1% yeast extract, 2% bactopeptone, 2% agar) at 4°C.

**Table 1.**
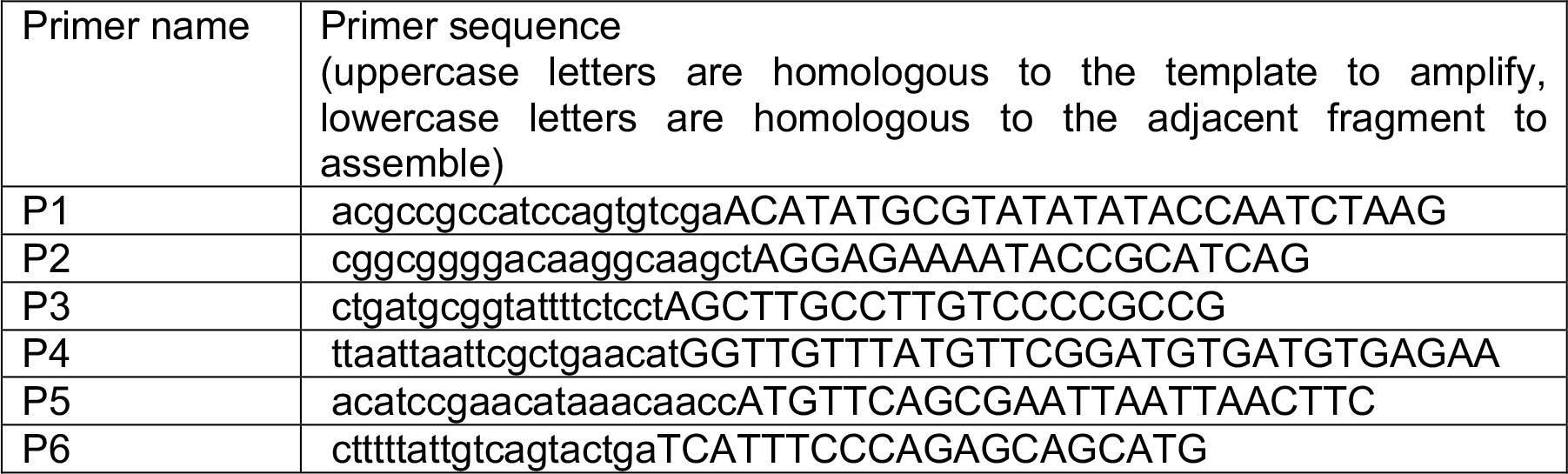
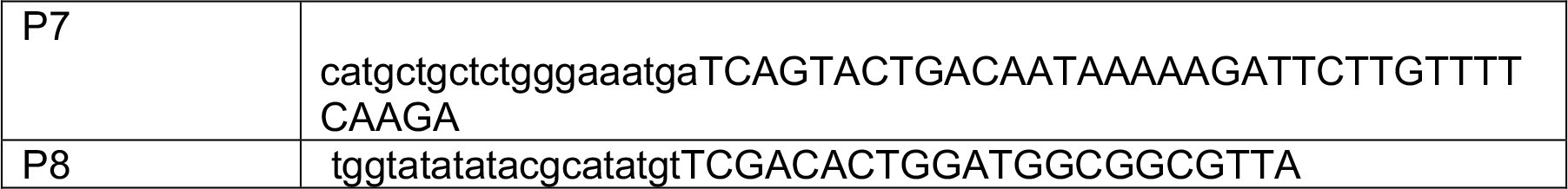
Primers used in this work.

### Fermentation conditions and H_2_S quantification

Fermentation experiments were conducted using synthetic must (SM), designed to mimic the characteristics of a natural grape must ^38^. It contained a 200 g/l equimolar glucose and fructose content,), and 200 mg/L assimilable nitrogen, 3.8 mg/l phytosterol, and 0.25 mg/l of Cu2+. The pH was adjusted to 3.3 with sodium hydroxide solution.

One single colony of each strain was grown in 5 ml of liquid YPD at 28°C for 24 hours and then diluted 100 times in SM. After 24 hours at 28°C, cells were counted with an electronic particle counter (Multisizer 3 counter; Beckman Coulter) and 250 mL of SM, supplemented with 60 mg/L of SO_2_ when the impact of sulfite was evaluated, were inoculated to 1 x 10^6^ cells/mL. Fermentations were carried out at 28°C, under permanent stirring (280 rpm) and they were followed daily by weight loss, until the theoretical percentage of sugar consumed reached 95% (87.4 g CO_2_/L produced). Total H_2_S produced during alcoholic fermentation was collected with a zinc-based trap system and quantified with sulfide specific fluorescent probe, as described before ^39^.

When the impact of the overexpression of *CUP1* was in study, SM was supplemented with Geneticin (G418 – Sigma A1720-5G) to maintain the plasmid allowing the overexpression itself. Suitable antibiotic concentrations were defined for each strain (100μg/mL for wine strains, 40 μg/mL for the oak one), to simultaneously allow the maintenance of the plasmid and a good fermentation rate, but prevent the growth of the sensitive strain (i.e. the wild-type strain without the plasmid).

When assessing the impact of copper concentration on H_2_S production, SM was supplemented with copper sulfate to reach 1 or 2 mg/L of copper; control copper concentration was 0.25 mg/L in all the experiments. More details about the experiments are given in the section “Experimental design and statistical analysis”.

### *CUP1* copy number evaluation

For most of the strains, *CUP1* copy number was estimated from their genome sequence, obtained from previous works or from sequencing performed in this study. To obtain the values, the median sequencing depth measured at SNPs encountered between coordinates 212500 and 213000, and between 214500 and 215000 on Chromosome VIII was divided by the median sequencing depth over the entire genome (excluding mitochondria and 2 microns). For Italian strains, *CUP1* copy number data had been already quantified by Real Time PCR ^18^.

### Genomic DNA extraction for sequencing

Genomic DNA was isolated from liquid yeast cultures in stationary phase, with a classical phenol-chloroform method, as described before ^40^, with an additional purification step based on the use of silica-coated magnetic beads (GMG-252-A-100mL – PerkinElmer), as follows. Cells were broken mechanically by shaking them in the presence of 600 μm diameter glass beads, lysis buffer (Tris 50 mM pH 8, EDTA 50 mM, NaCl 100 mM, Triton 2%, SDS 1.25%) and phenol chloroform isoamyl alcohol 25:24:1. DNA was precipitated with ispopropanol and ethanol, dried, resuspended in TE (Tris 10 mM, EDTA 1 mM) and treated with RNase A. Samples were mixed with the DNA absorption solution (for one sample: 50 uL 5 M NaCl, 15 ul magnetic beads (GMG-252-A-100mL – PerkinElmer), 250 uL 7.8 M guanidium chloride, 800 uL isopropanol), after which metal beads with DNA absorbed on their silica surface were recovered using the DynaMag™-2 Magnet tube holder (12321D-DynaMag-2 – Invitrogen) and washed twice with AMMLAV/E buffer (10 mM Tris pH 8, 0.1 mM EDTA, 60 mM potassium acetate, 65% ethanol) and twice with ethanol 75%. DNA was then desorbed and in aqueous solution.

DNA purity was checked from the 260_nm_/280_nm_ and 260_nm_/230_nm_ OD ratio measured with NanoDrop 1000 (ThermoScientific). The DNA was quantified by fluorescence using the QuantiFluor kit, dsDNA system (Promega) and then stored at -20°C.

### Genome Sequence and Analysis

DNA samples were processed to generate libraries of 500 bp inserts. After passing quality control, the libraries were sequenced with DNBseq technology using BGISEQ-500 platform, generating paired-end reads of 2 × 150 bp.

For each library, low-quality reads were processed and filtered using the FASTX Toolkit v0.0.13.2 and TRIMMOMATIC v0.36 ^41^ with the following parameters (LEADING:10 HEADCROP:5 SLIDINGWINDOW:4:15 MINLEN:50).

Reads were then mapped to the S288C reference genome with BWA v0.6.2 with default parameters ^42^ and genotyping made with samtools v1.11 to obtain a variant file including the sequencing depth of each variant position. Sequence positions were afterwards filtered for quality criteria: sufficient coverage position as well as genotyping and mapping quality (MQ >-20) were kept.

### Plasmid construction and yeast transformation

*CUP1* was inserted via Gibson assembly method ^43^ between TEF promoter and terminator in a high copy Yeast Episomal plasmid (YEp352), modified to confer geneticin resistance (YEp352-G418) to the host cell. In detail, the backbone was amplified with primers P1 and P2, designed to replace the original *URA3* copy of YEp with *CUP1*, since the strains used were not auxotrophic and the selection had been made by antibiotic. Therefore, the backbone contained a 2 um replication origin (multicopy), AmpR, ColE1, pPGK and G418 resistance cassette. *CUP1* was amplified from OakGri7_1, a strain previously sequenced by our laboratory ^10^, with a single metallothionein copy and the same sequence as laboratory reference strain S288C, used to design primers (P5 - P6). TEF promoter and terminator were amplified from pCfB2312 ^44^ with primers P3 - P4 and P7 - P8, respectively. Primer sequences are listed in Table 1.

Proper fragment insertion was verified by enzymatic digestion (NarI, ClaI, PacI - New England Biolabs). To assure that the phenotype was related to the overexpression of *CUP1*, a Yep352-G418 plasmid without *CUP1* was used as control. PCRs were performed with Phusion™ High-Fidelity DNA Polymerase and validated by gel electrophoresis. *Escherichia coli* strain DH5α was used to maintain and amplify the plasmid; cells were selected on LB medium with ampicillin (100 μg/ml) and grown at 37°C. Yeasts (Oak-Rom 3-2, LMD17 and L1374) were transformed with the lithium acetate method ^45^ and strains containing the recombinant plasmids were selected on YPD agar with 200 μg/mL geneticin (G418 – Sigma A1720-5G).

### Experimental design and statistical analyses

Experiment 1: impact of the origin of the isolate and sulfites on H_2_S production 33 strains were selected randomly from our laboratory collection (Supplementary table 1, dataset 1). Alcoholic fermentations were performed in absence or presence of SO_2_, in duplicate for each strain and each condition.

The factors accounting for the variation of H_2_S were analyzed with the following analysis of variance model:

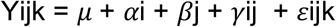

where Yijk is the H_2_S production, μ the overall grand mean, *α*i is the fixed strain group effect, *β*j is the fixed SO_2_ effect, γij is their interaction effect, and *ε*ijk the residual error.

The analysis of the residuals showed that three values were distant from the global distribution. Since results of the statistical analysis did not change after removing all the observations of the three outlier strains, the complete dataset was kept as the method is sufficiently robust to mild deviations.

Experiment 2: impact of copper content of the media on H_2_S production Fermentations were performed without SO_2_ in triplicate, for each strain (VL1 and LMD17) and each condition (0.25 - 1 and 2 mg/L of copper).

To evaluate the effect of copper and strains on H_2_S production, ANOVA was performed, after checking for the equality of variance with a Levene test. The most parsimonious model was kept after checking of the absence of interaction between strain and the copper content:

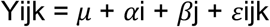

where Yijk is the H_2_S production, μ the overall grand mean, *α*i is the fixed strain effect, *β*j is the fixed copper effect and *ε*ijk the residual error.

Experiment 3: impact of *CUP1* copy number on H_2_S production To the strains evaluated in experiment 1, we added 18 wine strains (total strains analyzed = 55), some known to harbor a high number of copies of *CUP1*, and some commercial strains known to be high H_2_S producers (Supplementary table 1, dataset 2). Alcoholic fermentations were performed in absence of SO_2_, in duplicate for each strain.

Different polynomial models were used to describe the interaction between H_2_S production and *CUP1* copy number (first-, second- and third-degree polynomial models); ANOVA was used to assess the significance of these models.

Experiment 4: impact of the overexpression of *CUP1* on H_2_S production Fermentations were performed without SO_2_ and with standard copper content (0.25 mg/L) in triplicate, for each strain (OAK_ROM 1-3, LMD17 and L1374) and each condition (wild-type strain, strain with the empty vector, strain with the CUP1 overexpressing vector).

ANOVA was performed to test the effect of the genetic modification in each strain. The model used was:

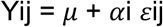

where Yij is the H_2_S production, μ the overall grand mean, *α*i is the fixed genetic modification effect, and *ε*ij the residual error.

For all the experiments, when the impact of one (or more) factor was significant, differences between modalities were evaluated by post-hoc testing (Tukey’s HSD multiple-comparison test, p < 0.05).

Statistical analyses were performed in the R environment (R version 4.0.2 (2020-06-22) _46_).

## Data availability statement

The datasets generated and analysed during the current study are available at Data Gouv (https://entrepot.recherche.data.gouv.fr/) with the following doi: https://doi.org/10.57745/5ECVDJ.

Genome sequence were deposited at EBI (https://www.ebi.ac.uk/) and corresponding accession numbers are given in Supplementary table 1 Strain-dataset.xlsx

## Acknowledgments

The authors thank Jessica Noble (Lallemand SAS), Prof Joseph Schacherer (University of Strasbourg) and Prof Viviana Corich (University of Padova) for providing the strains analyzed in this work.

All the authors are grateful to Thérèse Marlin for providing her expertise with the Gibson assembly strategy and for the molecular cloning of *CUP1*, to Dr Thibault Nidelet for the polynomial models and Dr Delphine Sicard for the statistical counseling and the proofreading of the manuscript.

## Author Contributions

BB and JLL conceived the study and supervised the work; IDG carried out the experiments and wrote the manuscript with support from JLL and BB; VG and IDG conceived the molecular biology experiments. All authors discussed the results and contributed to the final manuscript. All authors have read and agreed to the published version of the manuscript.

